# CCR2 antagonism identifies blood-borne monocytes as a target for prevention of cognitive deficits after seizures

**DOI:** 10.1101/2025.05.19.655010

**Authors:** Soheila Pourkhodadad, Wenyi Wang, Ray Dingledine, Nicholas H. Varvel

**Affiliations:** Department of Pharmacology and Chemical Biology, Emory University School of Medicine, Atlanta, GA

**Keywords:** Inflammation, chemokines, status epilepticus, disease prevention, epilepsy, cognition, co-morbidities

## Abstract

Seizure-associated cognitive co-morbidities can substantially reduce the quality of life in people with epilepsy. Neuroinflammation is an invariant feature of all chronic neurologic diseases, including epilepsy, and acute brain insults, including status epilepticus (SE). The generalized seizures of SE trigger a robust inflammatory response involving astrocytosis, erosion of the blood-brain barrier (BBB), activation of brain-resident microglia, and recruitment of blood-borne CCR2+ monocytes into the brain. We have shown that blocking monocyte recruitment into the brain via global *Ccr2* knockout or systemic CCR2 antagonism with a small molecule alleviates multiple deleterious SE-induced pathologies, including BBB damage, microgliosis, neuronal damage, and monocyte brain invasion in the days following pilocarpine-induced SE. This study aimed to determine if fleeting CCR2 antagonism improves SE-associated cognitive impairments in the long term. Here, we show that the brief antagonism of CCR2 eliminates the profound deficit in working memory in the Y-maze and retention memory in the novel object recognition test but does not attenuate anxiety-like behavior in the open field arena. Microgliosis and astrocytosis were observed in brain sections from SE mice after the behavioral tests, and CA1 hippocampal astrocytosis mildly correlated with performance in the Y-maze. Notably, SE mice exposed to the vehicle showed robust neuronal loss in the cortex and CA1 region of the hippocampus, and mice treated with the CCR2 antagonist showed less neurodegeneration in both the cortex and hippocampus. Our results indicate that monocyte brain infiltration after SE opens a window for preventing cognitive co-morbidities and neurodegeneration with an orally available CCR2 antagonist.

**Highlights:**

- CCR2 antagonism provides protection against seizure-associated cognitive deficits.
- A therapeutic window to alleviate SE-associated cognitive decline has been identified.
- Selective immune modulation might be used to treat epilepsy co-morbidities.

**Graphical abstract:** 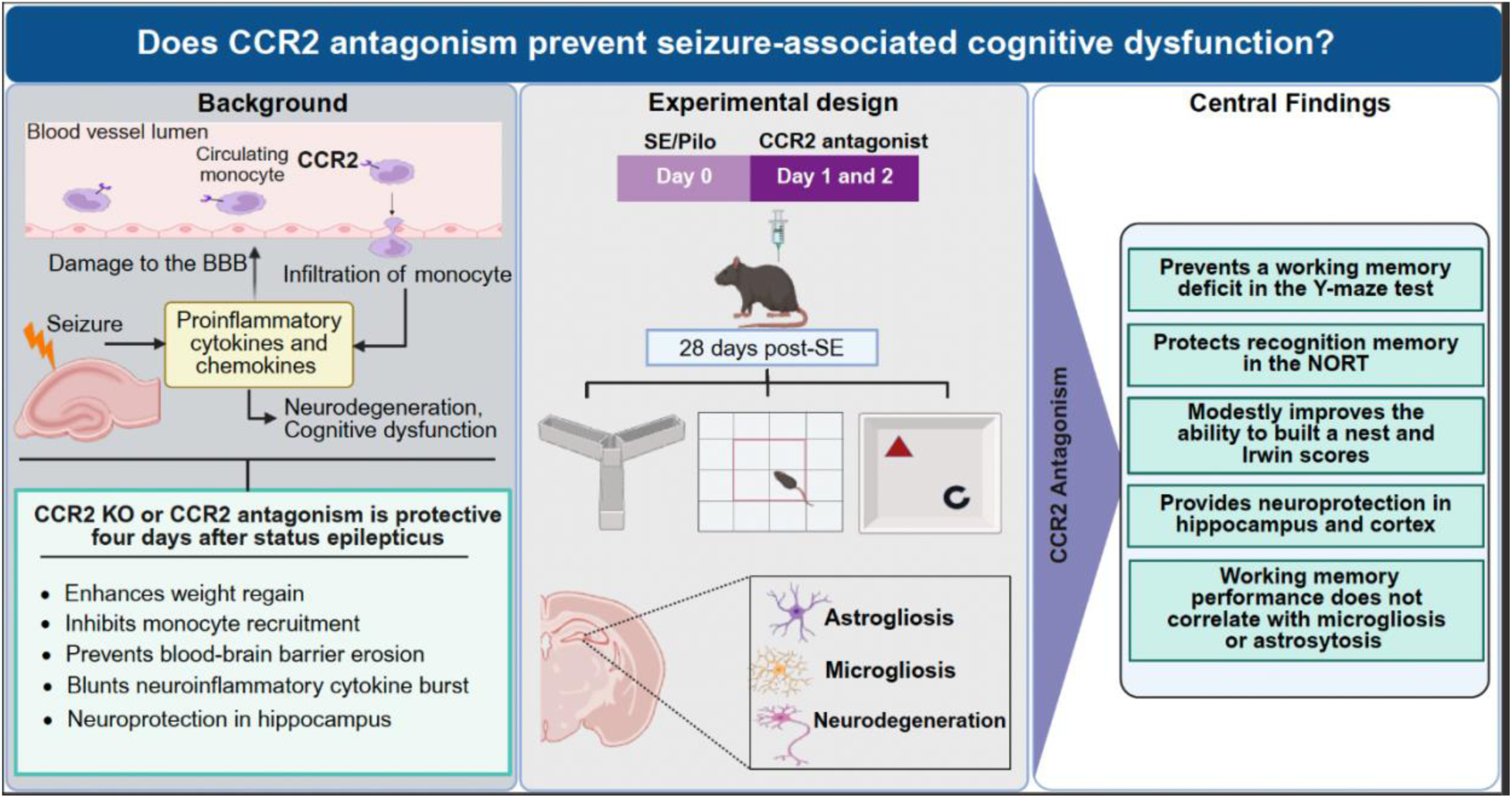

## Introduction

Epilepsy is a family of seizure disorders that affect over 3.4 million individuals in the United States and 65 million people worldwide. Epilepsy represents the fourth most prevalent neurologic disorder after stroke, Alzheimer’s disease, and migraine headaches. Epilepsy is frequently associated with memory impairments and psychiatric co-morbidities such as depression, anxiety, and mood disorders (Lu 2021, Mula 2022), which can significantly impact the quality of life and may occur independently of seizure activity. Among neurologic disorders, epilepsy accounts for the highest disability-adjusted quality life years lost (Beghi 2016). Strategies to modify or prevent epileptogenesis or seizure-associated co-morbidities are two of the arduous challenges in epilepsy care today (Galanopoulou 2016, Goldman 2016). Despite several notable recent clinical advances in treating epilepsy, such as neurostimulation, MRI-guided stereotactic laser surgery, and the introduction of new antiseizure drugs, one-third of patients cannot be managed by surgery or the existing drugs (Chen 2018, Beghi 2019). Moreover, long-term use of some antiepileptic drugs is associated with cognitive decline (Meador 2006, Park 2008).

The seizures of status epilepticus (SE) can cause memory deficits in affected patients that appear within weeks and can persist for a year or longer (Power 2018, Power 2018). In one report, over half of those experiencing convulsive SE lasting a median of 85 minutes had adverse cognitive outcomes such as the inability to return to work and live independently (Legriel 2010). Another study of ICU patients with refractory SE also reported poor long-term outcomes (Kantanen 2017). The incidence of SE in the USA is 18-41 per 100,000 people per year (Lu 2020). For those acquired epilepsies not caused by SE, cognitive deficits are often encountered at the time of diagnosis (Witt 2015). In rodent models of SE, cognitive and memory functions can be impaired prior to the development of spontaneous recurrent seizures, i.e., epilepsy (Hort 1999, Mohammad 2019). Preclinical and epidemiological observations support the idea that temporal lobe epilepsy and its cognitive co-morbidities, including memory impairments and anxiety disorders, may share similar pathological processes (Helmstaedter 2007, Bell 2011, Kleen 2012, Goldman 2016, Kanner 2016, Leeman-Markowski 2016). Sclerosis-associated neuronal death and/or neuroinflammation have been suggested as drivers of seizure-associated cognitive co-morbidities (Goldman 2016, Paudel 2018).

Nearly all acute brain injuries such as stroke and SE and chronic neurologic diseases such as Alzheimer’s disease (AD) and epilepsy feature an inflammatory component typified by a robust microglial response, reactive astrocytosis, erosion of the blood-brain barrier (BBB), and engagement of several broad inflammatory signaling cascades including cyclooxygenase-2 (COX-2) (Rojas 2014, Dingledine 2024, Heneka 2024). Infiltration of monocytes from the blood into the brain is a prominent feature of drug-resistant epilepsy patients, and patients who died shortly after a bout of SE (Broekaart 2018). We have shown a prominent role for blood-borne monocytes in the florid neuroinflammatory response to SE in mice. Limiting monocyte brain entry after SE is neuroprotective, quenches seizure-induced inflammatory burst in the hippocampus, prevents erosion of the BBB, limits microgliosis, enhances weight regain, and preserves nest-building capability in animal models (Varvel 2016, Tian 2017, Feng 2019, Aleman-Ruiz 2023). These salutary effects observed three to four days after SE onset suggest CCR2 as a target to alleviate the delayed adverse effects of SE. These observations encouraged us to hypothesize that blocking monocyte brain entry with a CCR2 antagonism might alleviate seizure-associated cognitive impairments in the weeks after SE onset.

The CCR2 antagonist INCB3344 is a potent, orally active human and mouse CCR2 antagonist with high selectivity. We now present in vivo pre-clinical efficacy data for INCB3344 with the systemic pilocarpine SE mouse model revealing alleviation of SE-associated behavioral deficits, cognitive impairments, and neuroprotection, all of which identify CCR2+ monocytes as a viable therapeutic target to lessen the adverse impact of SE. These findings highlight a deleterious role for peripheral CCR2+ monocytes in neurologic disease and support the continued development of strategies to limit monocyte brain recruitment in the clinic with the goal of relieving the debilitating co-morbidities of seizures and epilepsy.

## Materials and methods

### Ethics statement

Experimental procedures were performed in accordance with the National Institutes of Health Guide for the Care and Use of Laboratory Animals. The mouse utilization protocol (201900137) was reviewed and approved by the Institutional Animal Care and Use Committee of Emory University. The experiments reported here are in accordance with the ARRIVE guidelines.

### Animals

We predetermined the number of mice per group for behavioral studies to be 15-20, based on our previous experience (Levin 2012, Rojas 2016, Jiang 2020, Varvel 2023). Groups of 16-20 C57Bl/6 male mice (8-10 weeks old, average weight 25.1 gm ± 1.3 SD) were purchased directly from Charles River Laboratories and allowed to acclimate in Emory’s Animal Facility for one week prior to status epilepticus (SE) induction. Mice were kept at ambient temperature with a 12-hour light-dark cycle (7am – 7pm) and provided unlimited access to food and water. During the acclimation period, the mice were group housed (5 mice/cage).

### Pilocarpine-induced SE and drug administration for behavioral studies

The mice were numbered and batch-randomized to receive saline (10 ml/kg i.p.) or pilocarpine HCl (freshly prepared 270 mg/kg as free base, i.p., Sigma P6503) in saline. All mice first received terbutaline (2 mg/kg i.p., Sigma T2528) and methylscopolamine (2 mg/kg i.p., Sigma S8502) to alleviate respiratory and cardiovascular effects of pilocarpine, then 20 min later received saline or pilocarpine. Pilocarpine-treated mice experienced SE for 1 h, after which all mice (including saline-treated) were administered diazepam (10 mg/kg i.p. and 30 ml/kg in saline) to interrupt SE. During this period, seizures were classified as previously described (Varvel 2016, Varvel 2021) and scored every 5 minutes for seizure severity as follows: a score of 0 represents normal behavior (walking, exploring, sniffing, grooming); a score of 1 represents immobility, staring, jumpy or curled-up hunched posture; a score of 2 represents automatisms (repetitive blinking, chewing, head bobbing, vibrissae twitching, scratching, face-washing, “star-gazing”); a score of 3 represents partial or whole body clonus, occasional myoclonic jerks, shivering; a score of 4 represents tonic seizures that involve rearing and sometimes falling or “corkscrew” turning and flipping; a score of 5 (SE onset) represents nonintermittent seizure activity consisting of stage 3 or 4 repeatedly; a score of 6 represents wild running, bouncing, and tonic seizures; and a score of 7 represents death. Following SE, mice were fed moistened rodent chow, monitored daily, and injected with 5% (w/v) dextrose in lactated Ringer’s solution (0.5 ml i.p.) when necessary.

Twenty-four hours after SE onset, surviving mice were matched into two groups to approximately equalize the number of stage 4 seizures experienced during SE, then a coin flip determined whether a group would receive the CCR2 antagonist (100 mg/kg and 10 ml/kg, p.o.) 24 and 48 h after SE onset, or its vehicle (5% NMP, 5% Solutol HS-15, 30% PEG-400, and 60% citric acid (10 mM)). Drug administration design and dosing were based on our experience and pharmacokinetic data previously generated (Aleman-Ruiz 2023).

The experimenter was blinded as to whether each group received drug or vehicle and was not unblinded until the data had been analyzed. Mice were group housed in a warm (28°C) and humid environment for 2–3 days, after which they were separated into individual cages. A modified Irwin test (Varvel 2021, Varvel 2023) was performed to assess the health and well-being of mice after pilocarpine-induced SE. The test comprised eight parameters (ptosis, lacrimation, body posture, tremors, dragging body, hyperactivity or hypoactivity, coat appearance, splayed hind legs) that can be measured simply by experimenter observation. Each parameter was scored on a three-point scale (i.e., 0=normal, 1=mild to moderate impairment, and 2=severe impairment) with a total score ranging from 0 to 16. A total score of 0 as a sum of all eight parameters indicates a normal healthy mouse. A total score ranging from 1 to 16 indicates a mouse that is increasingly impaired. Each mouse’s ability to construct a nest from a cotton square was also assessed (Deacon 2006). The mice were monitored daily from days one to seven post-SE, then once a week.

### CCR2 antagonist, INCB3344

INCB3344 is a small molecule antagonist of the chemokine receptor, CCR2 (MedChemExpress, Monmouth Junction, NJ). INCB3344 has favorable selectivity and specificity to CCR2 (target IC50 of 10 ± 5 nM; selectivity 300-fold against chemokine receptors CCR5 and CCR1, two close homologues of CCR2, and >100-fold selective against a broader panel of G protein coupled receptors) (Brodmerkel 2005).

### Y-maze performance

Spatial working memory was assessed using the Y-maze task 28 days after SE. The apparatus consisted of three arms, designated X, Y, and Z, which extended from a central point at 120° angles (Figure 2A). The maze was elevated 40 cm above the floor and placed in a dimly lit room to minimize external distractions. Mice were individually introduced at the end of one arm and allowed to explore the maze freely for 5 minutes. During this period, the sequence of arm entries and the total number of entries were recorded. The spontaneous alternation rate was determined by calculating the ratio of consecutive entries into three different arms, expressed as a proportion of the total possible alternations, defined as the total number of arm entries - 2. A lower alternation rate was indicative of potential impairments in working memory, as it reflects the inability of the mouse to remember which arm was previously visited. As in previous studies (Varvel 2023), once a mouse committed to an arm it tended to travel all the way to the end, so there was rarely any question whether a mouse had entered an arm.

The number of arm entries in different mice ranged from 7 to 46 during the 5-minute assay. To circumvent this difference, and to restrict the assay to the period in which the mouse focused on exploring the maze rather than attempting to escape, we analyzed the first 20 arm choices of each mouse. Mice that failed to enter at least 20 arms were not included. Five out of 20 total saline mice were removed for this reason; two out of 20 pilocarpine mice treated with the CCR2 antagonist were removed, and no pilocarpine-vehicle mice were removed.

### Open field test

Anxiety-like behavior and locomotion in mice were assessed using the open field (OF) test 29 days after SE. The OF arena measured 40 × 40 × 30 cm; each mouse was individually placed in the center of the arena and allowed to freely explore for 10 minutes while their movements were video recorded using the ANY-maze video tracking software (insert company name and city). Anxiety-like behaviors were evaluated with the observer blinded to the treatment conditions by measuring the total time spent when a mouse was in the center zone with all four paws. The arena floor was cleaned with 70% ethanol after each session. Total immobile time in all zones and, separately, the center zone, distance traveled (in meters), and average speed (m/s) during the 10-minute assay were calculated.

### Novel object recognition test

The novel object recognition test (NORT) was performed on days 30-33 post-SE using the same apparatus as the open field test. To ensure cleanliness and remove odors from previous mice, both the arena and the objects were sanitized with 70% ethanol before and after each behavioral assessment. The NORT is commonly performed to assess memory function in laboratory animals, particularly their capacity to distinguish and recognize new objects. Four distinctly different and immovable objects of similar height and firmness were used to evaluate their discriminatory ability. The experiment consisted of three phases: habituation, training, and testing. In the habituation phase, each mouse was placed in the empty arena and allowed to freely explore for 10 minutes. Twenty-four hours later, the training phase followed during which each animal was exposed to two identical objects for 10 minutes, and the exploration time spent with the familiar objects was recorded. After an interval of 2 hours, the animals were tested again for 10 minutes, with one familiar object and one novel object. We designated this 2-hour test as the “short-term NORT” (Figure 4A). Two days later the habituation phase was repeated, then 24 hours later the NORT was performed. We designated this 24-hour test as the “long-term NORT” (Figure 5A). During the time between the short- and long-term NORT, six mice in the saline-treated group were erroneously sacrificed by laboratory staff. Thus, there are fewer saline mice in the “long-term NORT” experiment compared to the “short-term NORT”. Exploration time in both tests was defined as the duration during which the animal’s nose was within 2 cm of the object or when it made physical contact with the object. The discrimination index (DI) was then calculated to quantify the animal’s recognition ability using the following equation: DI= (time exploring novel object – time exploring familiar object)/ (time exploring novel object + time exploring familiar object).

### Tissue processing

Thirty-five days after SE, mice were anesthetized deeply with isoflurane, perfused through the heart with ice-cold PBS solution, and their brains rapidly removed from the cranium. The brain was immediately bisected through the midline, and the left hemisphere was fixed in 4% (w/v) paraformaldehyde for 24 hours at 4 C. After fixation the left hemisphere was cryoprotected in 30% (w/v) sucrose in PBS solution. The brains were then frozen in 2-methylbutane chilled with dry ice and sectioned coronally at 35 μm using a freezing/sliding microtome.

### Histology

Brain tissue sections, 35 μm in thickness, underwent immunofluorescence staining using a free-floating method. To reduce nonspecific binding, the sections were incubated in a blocking solution of 5% goat serum (Sigma, G9023) for 45 minutes. They were then exposed to primary antibodies overnight at 4°C: rabbit polyclonal anti-Iba1 (1:1000, Wako AB839504) to label microglia/macrophages and rat monoclonal anti-GFAP (1:500, Invitrogen cat:13-0300) to identify astroglia. The sections were subsequently incubated with secondary antibodies Alexa Fluor 488 goat anti-rabbit (1:400, Invitrogen, AB143165) and Alexa Fluor 594 goat anti-rat (1:400, Invitrogen, AB141374) for one hour at room temperature. After PBS washes, the stained sections were mounted onto slides, and a DAPI-containing antifade mounting medium (Vector H-1200) was applied to preserve fluorescence and counterstain nuclei.

For neuron counting, mounted 35 μm thick sections were rinsed in deionized water, incubated in 60°C cresyl violet solution for 90 seconds, briefly washed in deionized water and differentiated in 70% and 100% ethanol. Slides were then cleared in xylene for 2 x 5 minutes and cover-slipped with Permount mounting medium (Fischer Chemical).

### Microgliosis and astrocytosis

Fluorescent imaging was performed to analyze glial marker expression in the CA1, CA3, dentate gyrus, and cortex. Brain sections stained for Iba-1 and GFAP were captured using a Carl Zeiss Axio Observer A1 epifluorescence microscope with an AxioCam Mrc5 camera. To ensure optimal detection of differences between saline-treated controls and pilocarpine-induced SE mice receiving vehicle treatment, threshold levels were established for each marker. Iba-1 images were acquired with brightness set to −0.35, contrast at 11, and an exposure time of 750 ms, applying the “Moments” threshold in NIH ImageJ Fiji software. GFAP-stained images were captured with brightness at −0.93, contrast at 5.56, and an exposure time of 500 ms, using the “Li” threshold setting. These parameters were consistently applied across all images for comparability. The percentage of area occupied by each glial marker was determined based on thresholded regions, with three sections averaged per animal.

### Estimation of neuron numbers

Quantification of cortical neuron number was done on the left hemisphere using systemic random sampling of every 12^th^ section. For CA1 pyramidal neuron estimates, a set of every 12^th^ section throughout the entire hippocampus was investigated. Reliable anatomic boundaries for cortex (olfactory bulb/tubercle, corpus callosum/external capsule, pia mater, endopiriform nuclei, claustrum, amygdala, and subiculum; (Bondolfi 2002) and CA1 (CA1 stratum oriens, CA1 stratum radiatum, CA2, subiculum; (Calhoun 1998)) were defined according to the Franklin and Paxinos mouse brain atlas (Franklin 1996). Using the optical fractionator principle (West 1993), the cortex optical disector spacing was 600 x 600 μm with a 3-dimensional dissector 31.6 x 31.6 x 15 μm. The CA1 optical dissector spacing was 100 x 100 μm with a 3-dimensional dissector 15 x 15 x 15 μm. A 100x objective (Carl Zeiss, Oberkochen, Germany; 1.3 numerical aperture) was applied to count the topmost nucleolus coming into focus within each disector, yielding an average number of 463 ± 30.1 SD cortical and 177 ± 54.4 SD CA1 neurons per mouse. Tissue thickness was measured at every third disector location, using a focus drive with 0.1 μm accuracy. Stereologic analyses were conducted in part with the assistance of Stereo-Investigator software (MicroBrightField, Williston, VT, USA). The average measured section thickness and SD was 28.0± 0.2 μm for cortical and 26.9 ± 0.5 μm for CA1. Neuron numbers are reported for one hemisphere.

### Statistical analysis

All tests were performed in a blinded fashion with the experimenter unaware of the treatment group. The nonparametric Mann-Whitney test was used for comparing two groups in figure 1D. Fisher’s exact test was used in figure 2D. A repeated measures one-way ANOVA with Holm-Sidak correction for multiple comparisons was used for figures 1E and 1G. The non-parametric Friedman test with Dunn’s multiple comparisons was used for figure 1F. The one-way ANOVA with a post hoc Dunnet’s test was used to compare the saline no SE and the pilocarpine-antagonist groups to the pilocarpine-vehicle group in figures 1B, 2B, 2C, 3B, 3C, 3D, 3E, 3F, 7C, and 7E. To evaluate the data from the NORT a one sample t test was used to compare each experimental group in figures 4B, 4C, 5B, and 5C to a hypothetical DI value of 0. Pearson’s correlation was used in figures 6B, 6C, 7B, and 7D.

**Figure 1.**
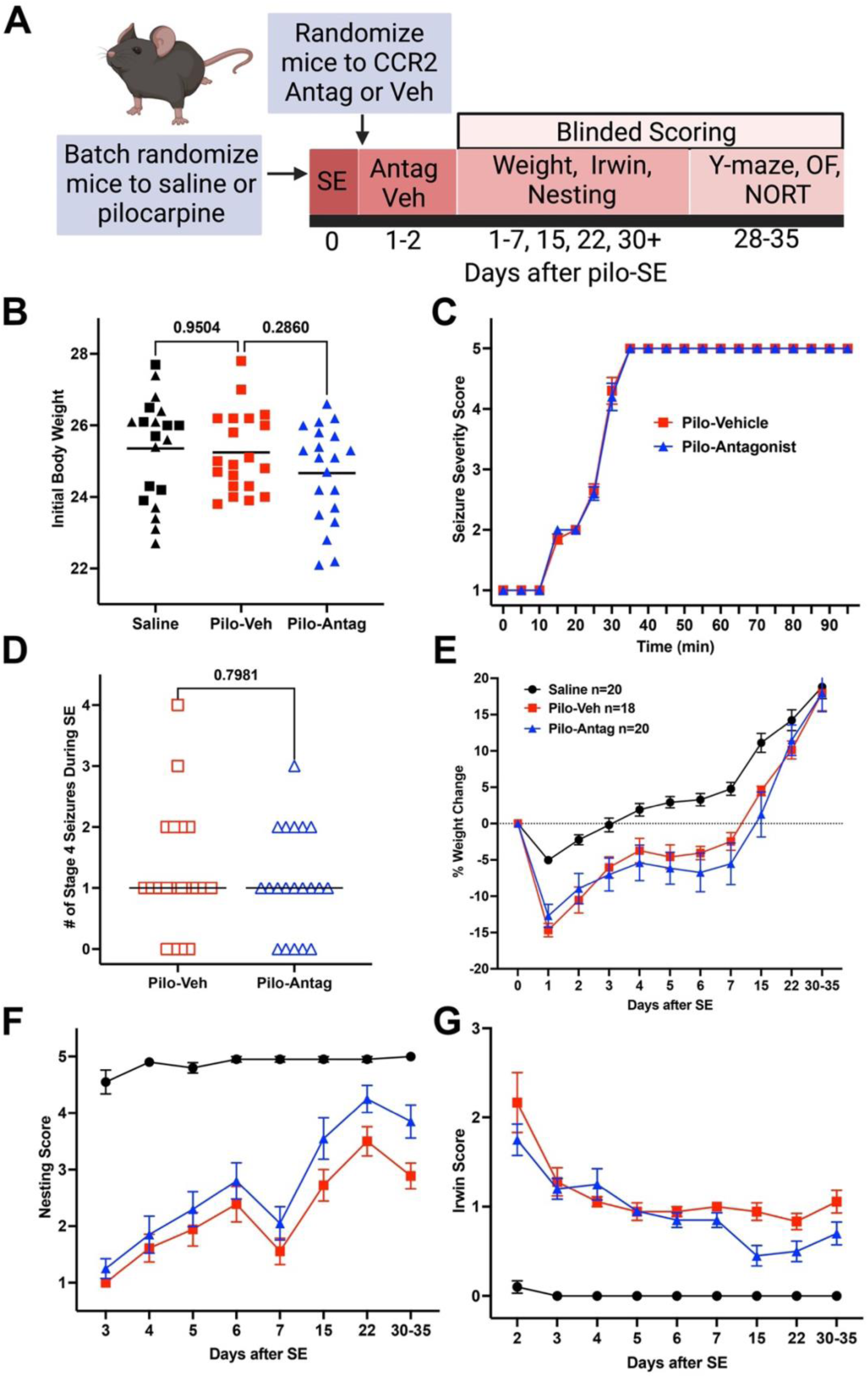
CCR2 antagonism modestly improves nesting capability and functional recovery scores. (A) Diagram of the experimental design showing the timeline of pilocarpine SE induction (day 0), randomization strategies prior to and after SE, CCR2 antagonist or vehicle dosing at 24 and 48 hours after SE onset, daily assessment of mouse weight, Irwin scores, and nesting ability, and behavioral tests for Y-maze, open field (OF), and novel object recognition tests (NORT). Created in BioRender.com. (B) Initial weight of all mice in the study (n=20/group). Saline-treated mice (no SE) were given either vehicle control (black squares) or CCR2 antagonist (black triangles). Data were compared with one-way ANOVA with Dunnett’s multiple comparisons test. Each symbol is from a different mouse, and horizontal line is at the mean. (C) Behavioral seizures were monitored for 90-95 minutes after pilocarpine administration. (D) Number of stage 4 seizures was similar between treatment groups, Mann-Whitney test. Each symbol is from a different mouse, and horizontal line is at the median. (E) Daily body weight for surviving mice in each experimental group. Significant differences were not observed between pilocarpine-vehicle and pilocarpine-antagonist groups, repeated measures one-way ANOVA with Holm-Sidak multiple comparisons test (p=0.0924). (F) Daily nest construction behavior of mice reveals a modestly improved ability to build a nest in the antagonist-treated group compared to vehicle-treated mice, Friedman test with Dunn’s multiple comparisons test (p=0.0455). (G) Daily Irwin scores show a mild improvement in functional recovery in mice treated with the antagonist compared to vehicle-treated mice, repeated measures one-way ANOVA with Holm-Sidak multiple comparisons test (p=0.0335). For 1E, 1F, and 1G, each symbol represents the group mean with SEM.

## Results

### CCR2 antagonism moderately alleviates adverse physiology after SE

Sixty-nine total male C57BL/6CR mice divided into four cohorts were batch-randomized to receive saline (n=22) or pilocarpine HCl (n=47), typically in a ~1:2 ratio to account for expected mortality in SE. All mice given pilocarpine HCl entered SE. Out of the 47 pilocarpine-treated mice that experienced SE, a total of six mice died in SE, and one mouse died within the next 18 hours before the first drug treatment. Two mice in the pilocarpine-vehicle group died after the second vehicle dose; no mice in the pilocarpine-antagonist group died. A diagram depicting our experimental design is shown (Figure 1A). The overall average body weight of the mice was 25.1 ± 0.2 grams, ranging from 22.1 to 27.7 grams. The body weights of the mice were not significantly different between each of the treatment groups. The saline-treated mice administered vehicle (Figure 1B, black squares) or CCR2 antagonist (black triangles) were grouped together for analysis in the current studies, because the CCR2 antagonist had no effect on any measured parameter in the control, saline-treated, mice. The temporal evolution of behavior seizure severity scores (Figure 1C) and number of stage 4 seizures by each mouse (Figure 1D) were similar between the pilocarpine-vehicle and pilocarpine-antagonist groups. Moreover, the mice were evenly distributed across the three treatment groups, which demonstrates that our batch randomization protocol was successful.

We monitored body weight, nest building capability, and sickness behaviors with a modified Irwin test for up to 35 days post-SE because our previous study revealed enhanced weight regain and modestly improved nest ability after CCR2 antagonism in the kainic acid (KA) SE model (Aleman-Ruiz 2023), and SE-induced inflammation is a strong driver of sickness behaviors (Varvel 2021). As expected, pilocarpine-treated mice lost ~12-15% of their initial body weight during the first 24 hours after SE onset. In contrast to our previous work, the CCR2 antagonist did not accelerate weight regain (Figure 1E) in the current study. However, SE mice given the CCR2 antagonist showed a modestly improved ability to build a nest based on the scoring described in Deacon RM (2006) (Figure 1F). A modified Irwin test wherein higher scores reflect more functional impairment was used to assess sickness behaviors. In general, modified Irwin scores were high 48 hours after SE and then declined toward normal regardless of treatment. SE mice given the CCR2 antagonist showed slightly lower modified Irwin scores over the course of our monitoring, suggesting less deterioration of normal physiological characteristics (Figure 1G). Relief from abnormal posture and enhanced grooming capabilities had a dominant contribution to the observed behavioral differences between the treatment groups.

### SE-induced working memory deficits are reduced by the CCR2 antagonist

Successive arm choices during free exploration of the 3-arm Y maze are viewed as a measure of spatial working memory because the innate curiously of rodents leads to exploration of the arm less recently visited (Hughes 2004). We observed that for many mice in the Y-maze (Figure 2A) their strategy shifts from exploration to escape as the 5-minute trial progresses. For this reason, we limit the alternation analysis to the first 20 arms entered. Five saline-treated and two pilocarpine-antagonist mice were removed from the analysis because they did not enter 20 arms.

**Figure 2.**
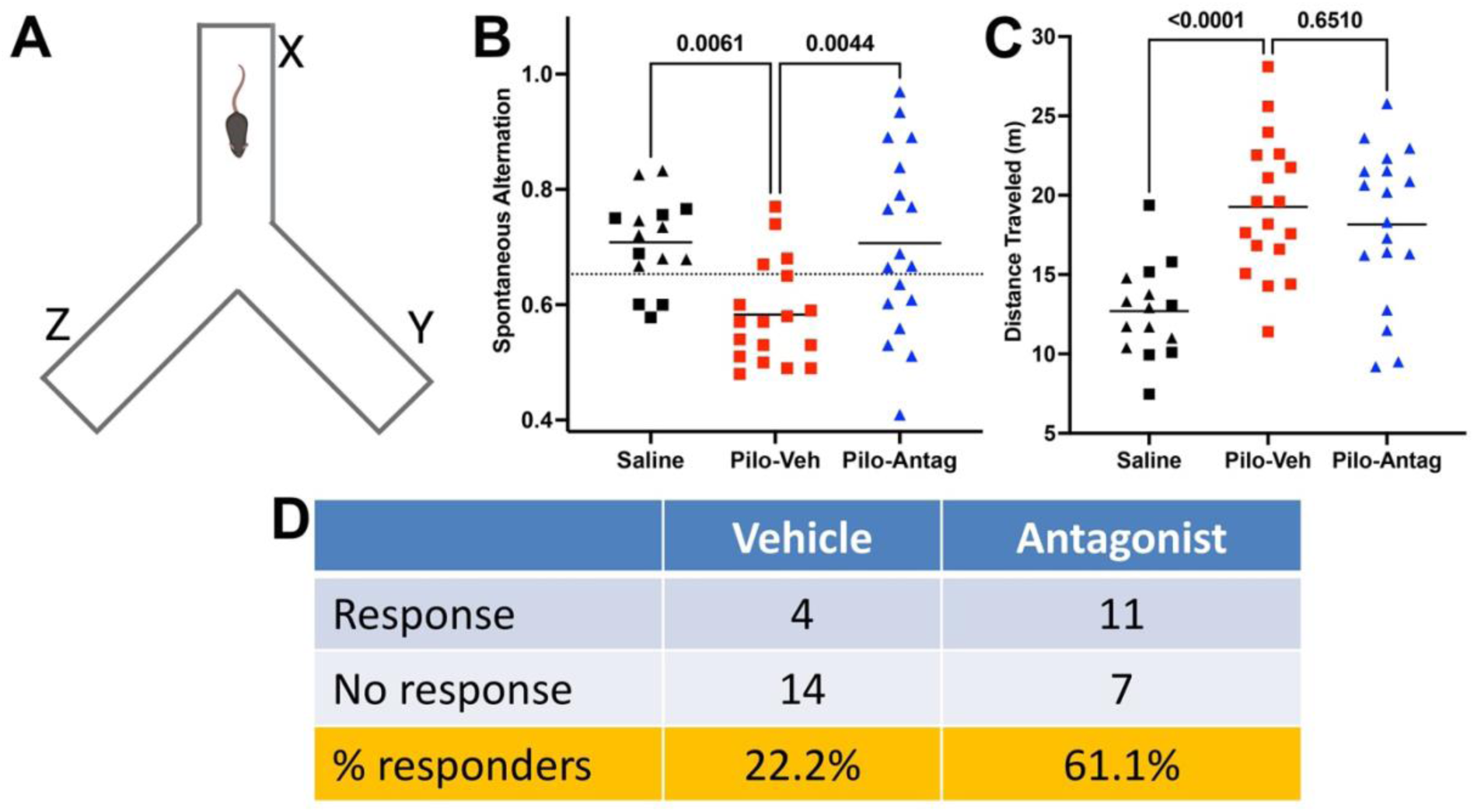
Brief exposure to the CCR2 antagonist improves working memory. (A) Diagram of Y-maze showing three arm choices, X, Y, and Z. Created in BioRender.com. (B) Mice were treated with pilocarpine or saline, then vehicle or CCR2 antagonist. 28 days after SE the mice were allowed to freely explore a Y-maze. After 20 arm choices the fraction of spontaneous arm choices that resulted in an alternation was calculated. Dotted line represents the value for a 50% improvement in mean spontaneous alternation (SA) score. (C) Distance traveled by each mouse during the Y-maze assay. Data were analyzed by one-way ANOVA with Dunnett’s multiple comparisons test in 2B and 2C, each symbol is from a different mouse, and the horizontal lines are at the mean value. (D) Table showing a significant improvement in SA score (defined by >50% improvement SA score) in antagonist-treated mice compared to vehicle-treated, Fisher’s exact test, p=0.0409.

The experiment (Figure 2B) was analyzed by one-way ANOVA. For the experiment, F (2, 48)=6.7945, p=0.0025. Control mice that received saline followed either by vehicle (black squares) or the antagonist (black triangles) alternated arm choices 70.8% of the time (Figure 2B), whereas pilocarpine-treated mice given vehicle performed at a degraded level (58.2%, red squares). However, SE mice with transient exposure to the CCR2 antagonist performed like control mice (70.7%, blue triangles). As an additional assessment of CCR2 antagonism on Y-maze performance, we asked what percent of SE mice given vehicle or antagonist showed >50% improvement in the assays. Eleven out of 18 (61.1%) pilocarpine mice treated with the antagonist showed >50% improvement, whereas only four out of 18 (22.2%) vehicle-treated mice showed improvement in the Y-maze (Figure 2D). Consistent with our previous findings (Varvel 2023), pilocarpine mice traveled a greater distance during the entire five-minute Y-maze test compared to saline-treated control mice (Figure 2C). Figure 2C also revealed that the enhanced performance of CCR2 antagonist-treated mice was not due to any difference in motor ability when compared to the pilocarpine mice subject to vehicle. For the SE mice, there was no correlation between distance traveled and y-maze performance (R^2^ =0.0044 and 0.069, not shown).

### CCR2 antagonism does not alleviate SE-induced anxiety-like behaviors

To assess potential differences in anxiety-like behavior and locomotion in the different groups of mice, we compared the percent time spent by each group in the center (anxiogenic) zone in the open field (OF) arena. Although it is difficult to ascribe emotions to rodents, less time spent in the anxiogenic zone is commonly interpreted as anxiety-like behavior. A diagram of the OF apparatus with the anxiogenic zone outlined in red taking up 37% of the entire arena is shown in Figure 3A. The 10-minute OF experiment was analyzed by one-way ANOVA. For the experiment (2,55) = 9.307, p=0.0003. Control mice that received saline followed by vehicle or CCR2 antagonist spent 13.8% of their time in the center anxiogenic zone (Figure 3B), whereas pilocarpine-treated mice given vehicle spent 7.6% of the time in the anxiogenic zone. The increased anxiety-like behavior was not relieved in the pilocarpine-treated mice given the CCR2 antagonist, and on average each group spent 1.4% and 0.6% of the assay time immobile in the center zone (Figure 3C). Administration of the CCR2 antagonist did not change immobility time in the angiogenic zone. In other movement parameters, including percent stationary time in all zones (Figure 3D), speed in the OF (Figure 3E), and total distance (Figure 3F), the pilocarpine-veh mice showed hyperactive behavior when compared to the saline-controls, and the hyperactivity was not alleviated in the CCR2 antagonist-treated group. Taken together, these findings reveal that pilocarpine increases locomotion and enhances anxiety-like behaviors in the OF test.

**Figure 3.**
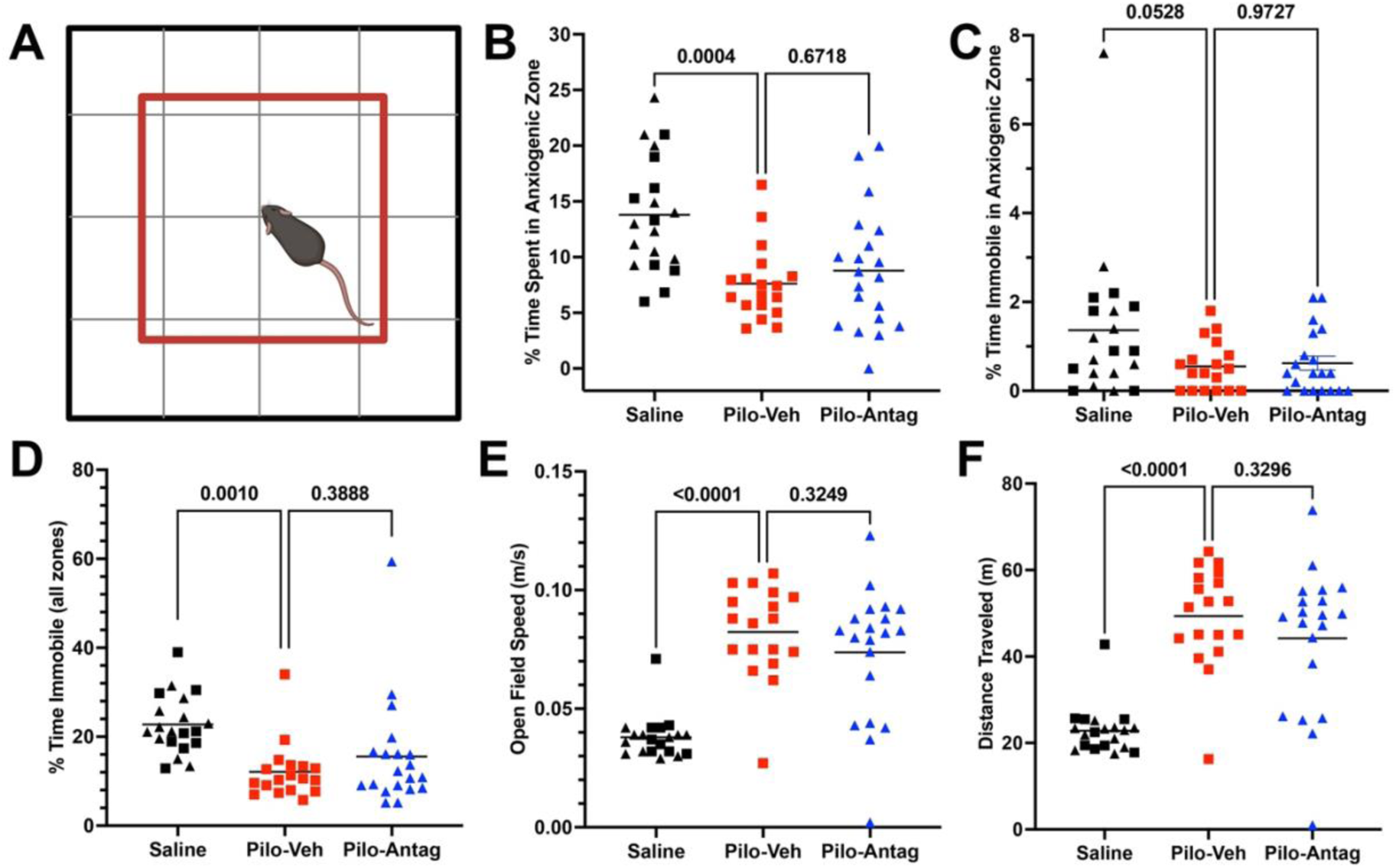
Anxiety-like behavior and locomotion are not alleviated by the CCR2 antagonist. (A) Diagram of the OF arena with anxiogenic (center) zone shown in red. The anxiogenic zone is 37% of the entire OF arena’s area. (B) Time spent in the anxiogenic zone was reduced in pilocarpine-vehicle mice compared to saline-treated no SE mice. One-way ANOVA F with Dunnett’s. (C) Percent time spent immobile in the anxiogenic zone was not significantly different between the groups. One-way ANOVA F(2, 55)=3.190, p=0.0489 with Dunnett’s. (D) Pilocarpine-treated SE mice administered spent less time immobile than saline-treated mice. One-way ANOVA F(2, 55)=7.195, p=0.0017 with Dunnett’s. (E) Pilocarpine-treated SE mice administered vehicle showed greater speed in the OF. One-way ANOVA F(2, 55)=26.31, p<0.0001 with Dunnett’s. (F) Distance traveled by the pilocarpine mice treated with vehicle was greater than saline-treated mice. One-way ANOVA F(2, 55)=25.94, p<0.0001 with Dunnett’s. Each symbol is from a different mouse, and the horizontal lines are at the mean value.

### Short-term recognition memory is not impaired after pilocarpine-induced SE

We next asked if CCR2 antagonism prevents the development of short-term memory impairments by comparing novel objection recognition performance among the three experimental groups using a discrimination index (DI). A DI score of zero indicates a mouse spends equal time with both objects. During the familiarization phase the saline group obtained an average DI score of −0.05, the pilo-vehicle group performed at 0.07, and the pilo-antagonist group at −0.01. Each group’s performance was not different from a DI score of zero by a 1-sample t-test, indicating a lack of preference for either identical object (Figure 4B). Two hours later a novel object was introduced into the arena (Figure 4A), and the mice were allowed to freely explore. The saline-treated mice spent more time exploring the novel object compared to the familiar object with an average DI of 0.17 (p=0.0046, 1-sample t-test compared to zero) (Figure 4C). Similarly, both pilocarpine groups also spent more time exploring the novel object with the pilo-vehicle group obtaining a DI score of 0.14 (p=0.0024) and pilo-antagonist group performing at DI=0.22 (p=0.0010). Taken together these data reveal the mice can discriminate a novel object from a familiar object, and short-term recognition memory is not impaired in pilocarpine mice under these experimental conditions.

**Figure 4.**
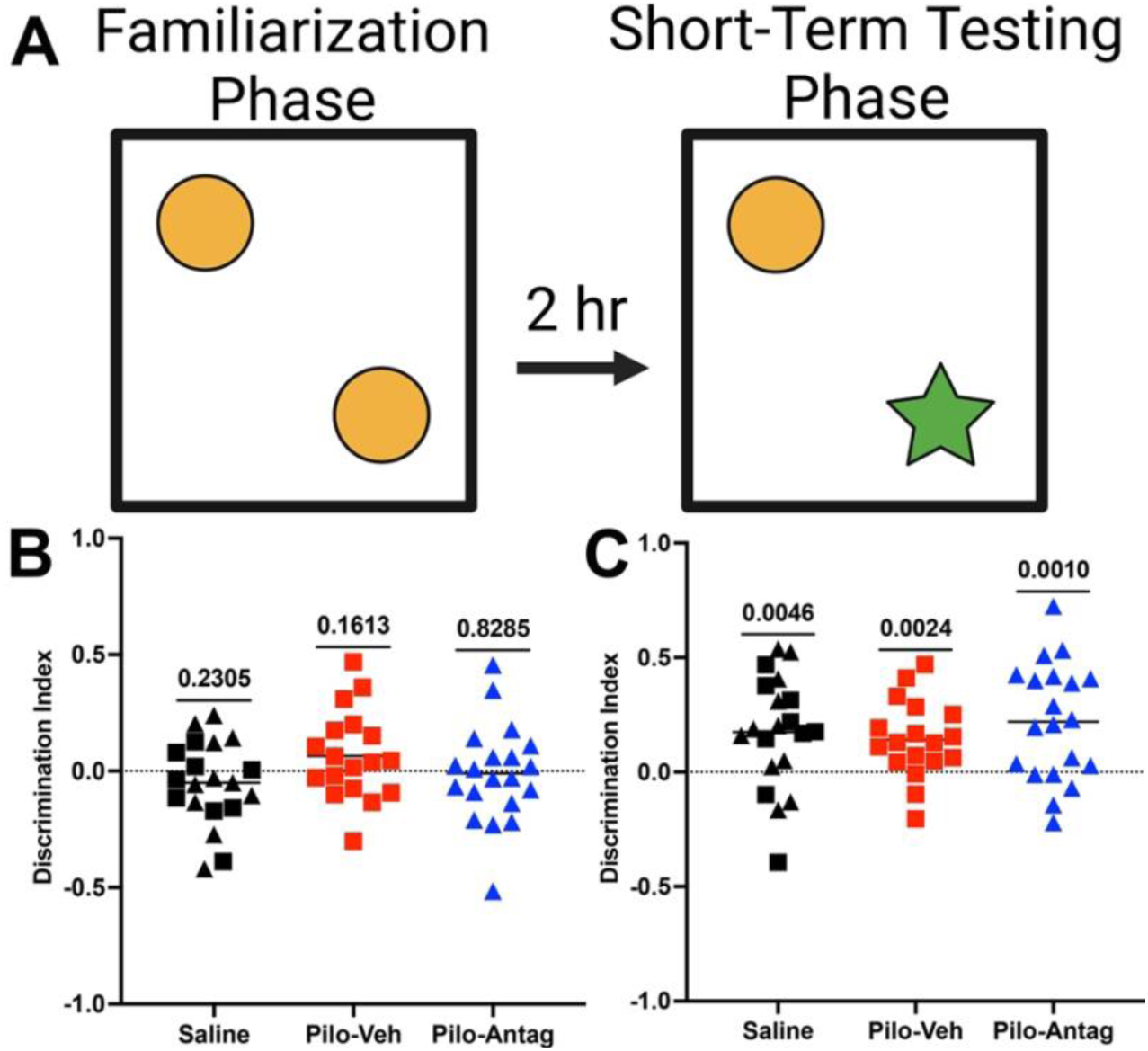
Pilocarpine SE mice do not show a deficit in short-term recognition memory in the NORT. (A) Diagram of the experimental design. Thirty days after SE, each mouse was allowed to freely explore two identical objects in the OF arena for 10 minutes. Two hours later one of the objects was replaced with a novel object, and the mice were allowed to explore for 10 minutes. A discrimination index (DI) was used as a measure of memory retention. The individual groups were compared to zero by a 1-sample t-test. (B) DI scores during the familiarization phase. (C) DI scores during the testing phase. Each symbol is from a different mouse. The horizontal dashed line at zero indicates the point at which there is no discrimination between the novel and familiar objects.

### CCR2 antagonism alleviates long-term recognition memory deficits

Given that the pilocarpine-vehicle group maintained the ability to identify a novel object two hours after the familiarization phase (Figure 4), we next asked if recognition memory deficits are encountered when the length of time between the familiarization phase and testing phase is extended to 24 hours (Figure 5A). During the familiarization phase, the saline (DI = −0.07), pilo-vehicle (DI = −0.03), and pilo-antagonist (DI = −0.04) groups all spent equal times exploring the two identical objects as each group’s score was not different from DI = 0 by a 1-sample t-test (Figure 5B). Twenty-four hours after the familiarization phase a different novel object from the short term test was placed in the arena, and the mice were allowed to freely explore for 10 minutes (Figure 5A). Both the saline (DI = 0.18, p=.0032, 1-sample t test) and pilo-antagonist (DI = 0.22, p = 0.0013, 1-sample t test) groups spent more time exploring the novel object, indicating that SE mice treated with the CCR2 antagonist were able to differentiate between novel and familiar objects after the 24-hour interval between exploration sessions. In contrast, the pilo-vehicle group (DI = 0.05, p=0.3782, 1-sample test) spent equal time exploring both the novel and familiar objects. These findings suggest that CCR2 antagonism alleviates SE-induced long-term recognition memory deficits.

**Figure 5.**
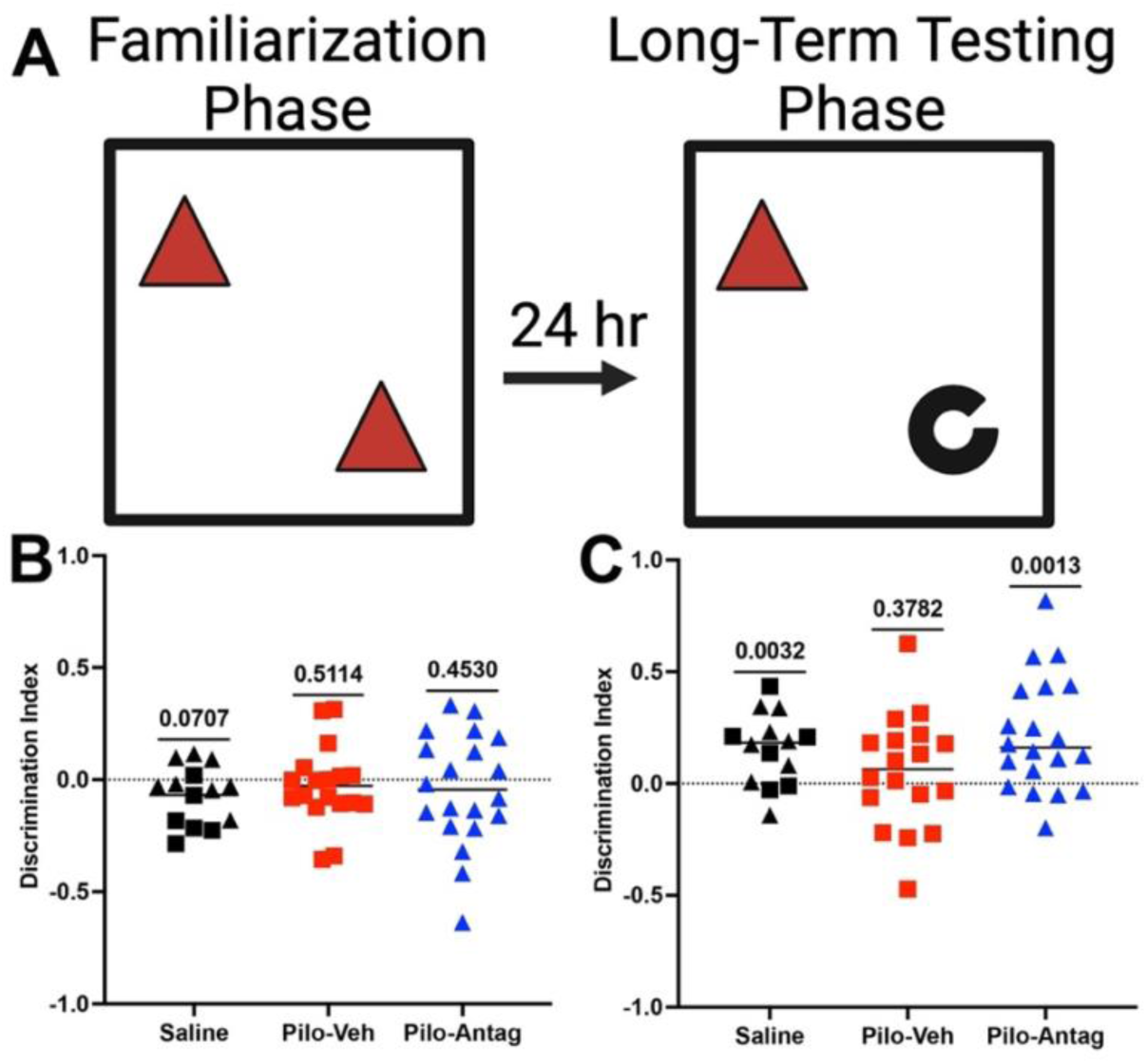
CCR2 antagonism alleviates deficit in long-term recognition memory in the NORT. (A) Diagram of the experimental design. Thirty-three days after SE, each mouse was allowed to freely explore two identical objects (different from short-term NORT) in the OF arena for 10 minutes. Twenty-four hours later one of the objects was replaced with a novel object, and the mice were allowed to explore for 10 minutes. A discrimination index (DI) was used as a measure of memory retention. The individual groups were compared to zero by a 1-sample t-test. (B) DI scores during the familiarization phase. (C) DI scores during the testing phase. Each symbol is from a different mouse. The horizontal dashed line at zero indicates the point at which there is no discrimination between the novel and familiar objects.

#### Sustained astrocytosis and Microgliosis do not correlate with Y-maze performance

SE causes robust neuroinflammation that is alleviated by CCR2 antagonism and in *Ccr2* knockout mice in the days following SE (Varvel 2016, Aleman-Ruiz 2023). Moreover, inhibition of the inflammatory prostanoid EP2 receptor after SE alleviates post-SE working memory deficits and suppresses microgliosis, but not astrocytosis or neuronal damage (Varvel 2023). Based on these data, we examined the correlation between Y-maze performance and astrocytosis and microgliosis by selecting the three best and three worst performers in each of our three experimental groups (saline control, pilo-veh, and pilo-antag). In this experiment brain tissue sections from 18 mice were subject to immunohistochemistry for GFAP (astrocytes) and Iba1 (microglia and monocyte-derived macrophages). Images were acquired from four brain regions in each mouse. Representative GFAP images are shown for CA1, with all acquisition settings held constant for all the brain regions and mice (Figure 6A). Figure 6B shows the correlation between SA score and the % area covered by GFAP staining in all four brain regions. Pearson r revealed mild inverse correlation between astrocytosis in the CA1 hippocampus and SA score. No correlation was found between microgliosis and SA score (Figure 6C).

**Figure 6.**
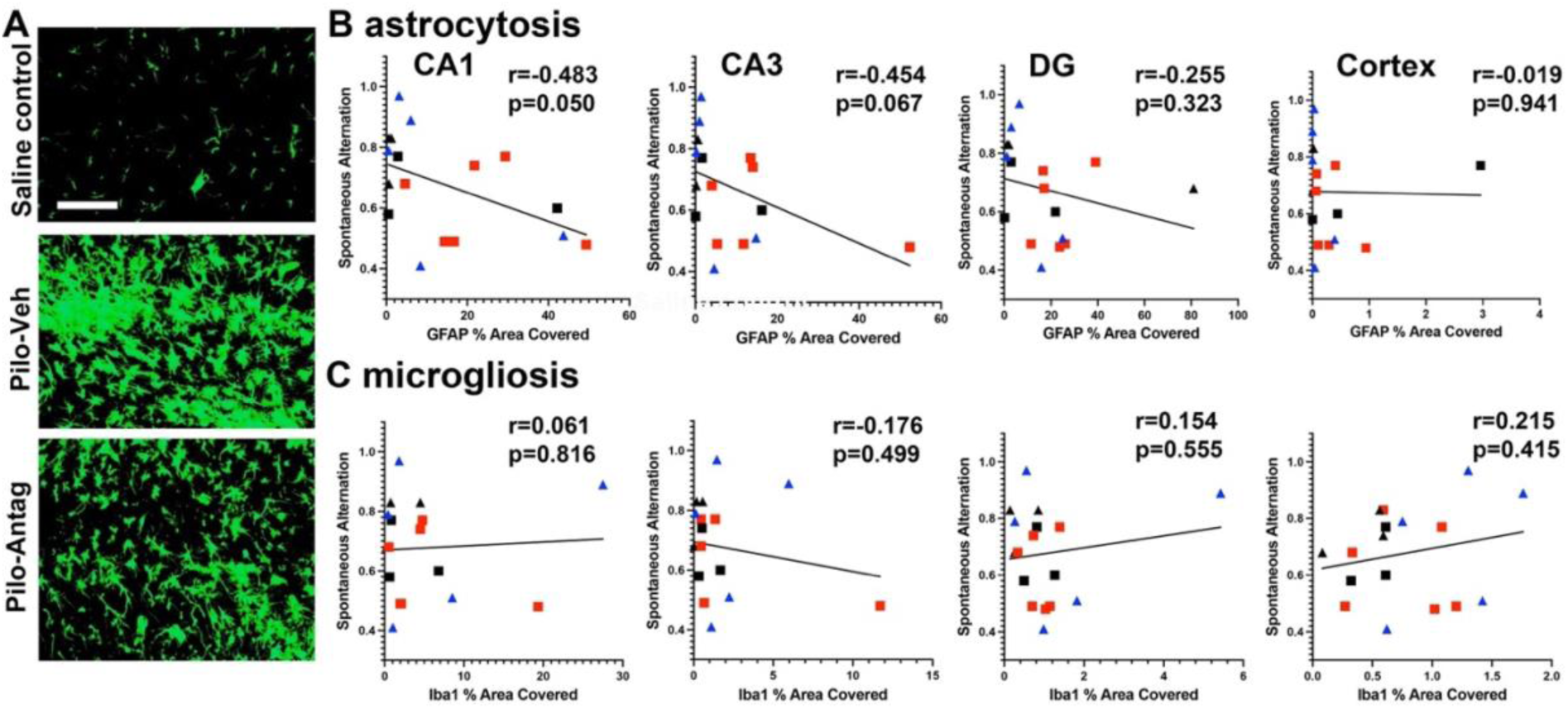
No correlation between inflammatory gliosis and spontaneous alternation scores in the Y-maze. (A) Representative images show positive GFAP immunostaining (green) as astrocytes in the hippocampal CA1 region. Thirty-five days after SE, astrocytosis is obvious in veh- and antagonist-treated SE mice compared with saline group. Scale bar 100 μm. (B) Pearson’s correlation coefficient analysis was performed to examine the relationship between SA score in the Y maze and brain region covered by GFAP immunostaining (n=18). (C) Pearson’s correlation coefficient analysis was performed to examine the relationship between SA score in the Y maze and brain region covered by Iba1 immunostaining (n=18).

#### CCR2 antagonism is neuroprotective

Neuronal loss is evident after SE. The same 18 mice selected for correlation analysis between gliosis and performance in the Y-maze were stained with cresyl violet to stain Nissl substance in the surviving neurons. Total numbers of cortical and CA1 hippocampal neurons were estimated using unbiased stereology and the optical fractionator method. Representative staining in the CA1 region is shown (Figure 7A). There was no correlation between Y maze performance and the remaining neurons in the CA1 hippocampus (Figure 7B) or the cortex (Figure 7D). The remaining neurons in the three groups were then compared using ANOVA with Dunnett’s. For the CA1 hippocampus, F(2,15) = 85.25, p<0.0001. The pilocarpine-veh group showed a 52.6% loss of CA1 neurons compared to the saline group, and the percent loss in the pilocarpine-antag mice was 19.4%. For the cortex, F(2,15) = 37.14, p<0.0001. The pilocarpine-veh group had a 13.2% loss compared to the saline group, and the pilocarpine-antag group showed a 5.6% loss.

**Figure 7.**
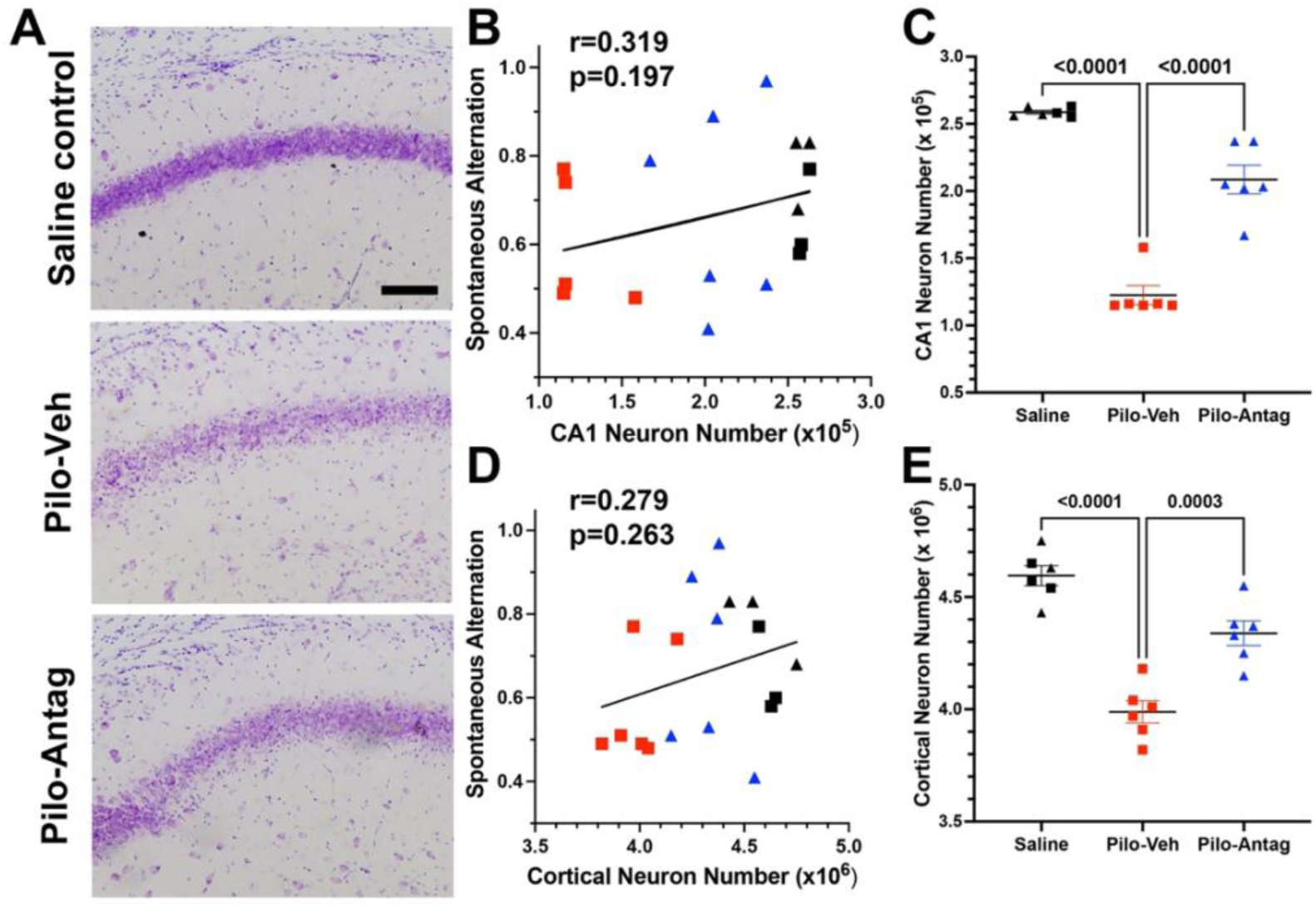
No correlation between neuronal numbers and spontaneous alternation scores in the Y-maze, but neuroprotection is evident. (A) Representative images show positive cresyl violet staining in the hippocampal CA1 region. Thirty-five days after SE, neuronal loss and layer dispersion is obvious in vehicle- and antagonist-treated SE mice compared with saline group. Scale bar 50 μm. (B) Pearson’s correlation coefficient analysis was performed to examine the relationship between SA score in the Y maze and CA1 neuronal number (n=18). (C) ANOVA with Dunnett’s showed a significant CA1 neuron loss in the pilocarpine-veh group and rescue in the pilocarpine-antag group. (D) Pearson’s correlation coefficient analysis was performed to examine the relationship between SA score in the Y maze and cortical neurons. (E) ANOVA with Dunnett’s showed a significant neuronal loss in the cortex in the pilocarpine-veh group and rescue in the pilocarpine-antag group.

## Discussion

Our study was designed to evaluate the effects of pharmacological inhibition of the inflammatory CCR2 receptor on seizure-associated cognitive co-morbidities. Cognitive impairments caused by SE and epilepsy are primarily treated with rehabilitation and are usually refractory to available pharmaceutical treatments or rehabilitation therapy (Brooks-Kayal 2013). Thus, our aim was to identify pharmacological targets and strategies for seizure-associated cognitive impairments. We found that systemic and fleeting treatment with a CCR2 antagonist after pilocarpine-induced SE alleviated a pronounced deficit in working memory in the Y-maze and recognition memory in the NORT when performed 24 hours after the familiarization phase. Notably, all three experimental groups showed discrimination indexes greater than zero when the NORT was performed two hours after the familiarization phase, indicating that short-term recognition memory was not impaired by pilocarpine SE. SE mice subjected to vehicle showed more anxiety-like behavior than the saline-treated control mice, but this condition was not rescued by CCR2 antagonism suggesting that the antagonist is not an anxiolytic. Whereas working memory performance in the Y-maze was not correlated with microgliosis or neuronal numbers in the cortex and CA1 hippocampus, there was a mild inverse correlation between astrocytosis in the CA1 hippocampus. Pilocarpine SE resulted in profound neuronal cell loss in the cortex and CA1 hippocampus, and the neuronal loss was attenuated in mice subject to the CCR2 antagonist.

The design and rationale for this study was based on previous findings wherein we and others first reported the involvement of CCR2-expressing monocytes in the neuroinflammatory response to both systemic kainic acid- and pilocarpine-induced SE (Varvel 2016, Tian 2017). The chemokine CCL2 is the principal agonist for Ccr2. In mice the mRNA levels of the key inflammatory mediators, Cox-2, IL-1β, and CCL2 are elevated within 30 minutes of SE onset, and microglial activation is evident in brain tissue sections 24 hours post-SE (Jiang 2015, Varvel 2016). Monocyte infiltration into the brain follows, starting between one and three days after SE onset (Varvel 2016). This delayed infiltration opens a clinically relevant therapeutic opportunity to target CCR2-expressing monocytes. Global *Ccr2* knockout reduces monocyte brain infiltration, accelerates weight regain, blunts hippocampal inflammatory burst, rescues erosion of the BBB, and is neuroprotective. The CCR2 antagonist, INCB3344, is orally active with acceptable plasma half-life of ~1.6 hr and shows brain to plasma ratio of 0.02-0.15 (Brodmerkel 2005, Xue 2010, Aleman-Ruiz 2023). Given our pharmacokinetic data showing robust bioavailability of INCB3344 in plasma, we calculate that a single 100 mg/kg dose (p.o.) of INCB3344 will provide at least 75% inhibition of CCR2 for 22 hours {Aleman-Ruiz, 2023 #5302}. Importantly, antagonizing CCR2 with INCB3344 after SE during peak monocyte brain invasion completely recapitulates the salutary consequences of global *Ccr2* deletion (Aleman-Ruiz 2023), confirming that the consequences of the *Ccr2* knockout were not a developmental effect. We have now extended our previous findings to include alleviation of seizure-related comorbidities as a beneficial consequence of CCR2 inhibition.

Interestingly, our group has shown that systemic antagonism of the prostanoid EP2 receptor with a brain-permeant small molecule also alleviates SE-induced neuropathology, blunts inflammation, and rescues a profound memory deficit after SE (Jiang 2012, Rojas 2015, Rojas 2021, Varvel 2021, Varvel 2023). Pharmacologic block of EP2 also prevents monocyte entry into the brain after SE (Varvel 2021). Brain-resident microglia express EP2, and expression is increased after SE (Quan 2013, Fu 2015, Varvel 2021). However, in contrast to microglia maintained in culture, freshly isolated microglia from the SE brain express a low level of EP2 mRNA, and blood-borne monocytes express nearly 100-fold higher levels of EP2 mRNA than microglia (Varvel 2021). The origin of brain-invading monocytes remains uncertain. It is possible the monocytes originate from the bone marrow or from limited privileged recruitment from restricted marrow passages in the skull (Cugurra 2021) or the spleen. These intriguing ideas are a question for additional studies.

The present findings implicate brain-invading monocytes as a driver of cognitive impairments after SE. Unanswered is whether CCR2 antagonism also prevents or attenuates recurrent spontaneous seizures in the chronic epileptic phase. Insights into the role of monocytes in seizures comes from the Theiler’s murine encephalomyelitis virus (TMEV) mouse model. Monocytes invade the brain after TMEV infection (DePaula-Silva 2018) and express high levels of the pro-inflammatory cytokine, IL-6 (Cusick 2013). Depleting monocytes and macrophages with liposome-encapsulated clodronate prior to TMEV infection reduces seizure frequency (Waltl 2018). However, limiting monocyte recruitment via *Ccr2* knockout did not prevent seizure development (Kaufer 2018). The reasons for these discordant findings are currently unclear. One possibility is that activation of peripheral immune cells and subsequent release of inflammatory mediators provoke seizures independent of monocyte migration into the brain; i.e., the involvement of CCR2 could be context dependent. Indeed, *Ccr2* genetic ablation or pharmacologic block did not mitigate the intense damage and neuroinflammation observed in a pediatric model of traumatic brain injury (Sharma 2023). Interestingly, peripheral inhibition of EP2 receptors with a brain impermeant antagonist quenched age-related hippocampal inflammation and restored memory deficits (Minhas 2021). These findings raise the intriguing possibility that peripheral, rather than central, inflammation might be a viable therapeutic target to alleviate neurologic disease. The benefit is that therapeutics would not have to be brain permeant to be effective.

The basis for cognitive impairments after seizures has remained obscure. SE-induced neuroinflammation, aberrant neurogenesis, and neurodegeneration are possible causative drivers (Parent 2002, Scharfman 2007, Parent 2008, Dingledine 2014, Rojas 2021). In support of neuroinflammation as a driver of cognitive deficits, febrile seizures can cause learning and memory deficits without overt neurodegeneration but provoke neuroinflammation (Dube 2009, Brennan 2021). Moreover, systemic EP2 antagonism after SE blunts neuroinflammation and rescues performance in the Y-maze but is not neuroprotective (Varvel 2023). Finally, in a peripheral inflammatory sepsis model with no neuronal degeneration, block of EP2 activation attenuates brain inflammation and preserves cognitive function (Jiang 2020). Limiting monocyte recruitment to the brain by targeting CCR2 also preserves cognition in sepsis models (Andonegui 2018). In the current study, markers of astrocytosis and microgliosis and neuronal cell number did not correlate strongly with Y-maze performance. However, Ccr2 antagonism was neuroprotective and rescued a profound memory impairment in the Y-maze. The common finding in these studies is that antagonizing EP2 or CCR2 prevents monocyte recruitment to the brain after inflammation provoking insults and alleviates cognitive deficits. If monocyte-induced brain inflammation is a driver of cognitive impairments, rather than neurodegeneration, then therapeutic reversal is practical.

Our study provides compelling evidence that transient pharmacological inhibition of the CCR2 receptor after SE alleviates several key seizure-associated cognitive impairments. By targeting monocyte infiltration during a therapeutically relevant window, CCR2 antagonism not only preserved recognition and working memory but also conferred neuroprotection. These findings strengthen the concept that brain-invading monocytes are critical drivers of SE-induced neuropathology and cognitive dysfunction. Together with previous work on EP2 receptor antagonism, our results suggest that targeting peripheral immune cell recruitment and activation could be a promising pharmacological strategy for treating cognitive comorbidities associated with status epilepticus in adults.

## Author contributions

N.H.V. conceived the idea. SP, WW, RD, and NHV preformed experiments. SP, WW, RD, and NHV analyzed and interpreted the results. SP, RD, and NHV wrote the manuscript. All the authors read and approved the final manuscript.

## References

Aleman-Ruiz, C., W. Wang, R. Dingledine and N. H. Varvel (2023). “Pharmacological inhibition of the inflammatory receptor CCR2 relieves the early deleterious consequences of status epilepticus.” Sci Rep 13(1): 5651.

Andonegui, G., E. L. Zelinski, C. L. Schubert, D. Knight, L. A. Craig, B. W. Winston, S. C. Spanswick, B. Petri, C. N. Jenne, J. C. Sutherland, R. Nguyen, N. Jayawardena, M. M. Kelly, C. J. Doig, R. J. Sutherland and P. Kubes (2018). “Targeting inflammatory monocytes in sepsis-associated encephalopathy and long-term cognitive impairment.” JCI Insight 3(9).

Beghi, E. (2016). “Addressing the burden of epilepsy: Many unmet needs.” Pharmacol Res 107: 79–84.

Beghi, E. and G. Giussani (2019). “Treatment of epilepsy in light of the most recent advances.” Lancet Neurol 18(1): 7–8.

Bell, B., J. J. Lin, M. Seidenberg and B. Hermann (2011). “The neurobiology of cognitive disorders in temporal lobe epilepsy.” Nat Rev Neurol 7(3): 154–164.

Bondolfi, L., M. Calhoun, F. Ermini, H. G. Kuhn, K. H. Wiederhold, L. Walker, M. Staufenbiel and M. Jucker (2002). “Amyloid-associated neuron loss and gliogenesis in the neocortex of amyloid precursor protein transgenic mice.” J Neurosci 22(2): 515–522.

Brennan, G. P., M. M. Garcia-Curran, K. P. Patterson, R. Luo and T. Z. Baram (2021). “Multiple Disruptions of Glial-Neuronal Networks in Epileptogenesis That Follows Prolonged Febrile Seizures.” Front Neurol 12: 615802.

Brodmerkel, C. M., R. Huber, M. Covington, S. Diamond, L. Hall, R. Collins, L. Leffet, K. Gallagher, P. Feldman, P. Collier, M. Stow, X. Gu, F. Baribaud, N. Shin, B. Thomas, T. Burn, G. Hollis, S. Yeleswaram, K. Solomon, S. Friedman, A. Wang, C. B. Xue, R. C. Newton, P. Scherle and K. Vaddi (2005). “Discovery and pharmacological characterization of a novel rodent-active CCR2 antagonist, INCB3344.” J Immunol 175(8): 5370–5378.

Broekaart, D. W. M., J. J. Anink, J. C. Baayen, S. Idema, H. E. de Vries, E. Aronica, J. A. Gorter and E. A. van Vliet (2018). “Activation of the innate immune system is evident throughout epileptogenesis and is associated with blood-brain barrier dysfunction and seizure progression.” Epilepsia 59(10): 1931–1944.

Brooks-Kayal, A. R., K. G. Bath, A. T. Berg, A. S. Galanopoulou, G. L. Holmes, F. E. Jensen, A. M. Kanner, T. J. O’Brien, V. H. Whittemore, M. R. Winawer, M. Patel and H. E. Scharfman (2013). “Issues related to symptomatic and disease-modifying treatments affecting cognitive and neuropsychiatric comorbidities of epilepsy.” Epilepsia 54 Suppl 4(0 4): 44–60.

Calhoun, M. E., K. H. Wiederhold, D. Abramowski, A. L. Phinney, A. Probst, C. Sturchler-Pierrat, M. Staufenbiel, B. Sommer and M. Jucker (1998). “Neuron loss in APP transgenic mice” Nature 395(6704): 755–756.

Chen, Z., M. J. Brodie, D. Liew and P. Kwan (2018). “Treatment Outcomes in Patients With Newly Diagnosed Epilepsy Treated With Established and New Antiepileptic Drugs: A 30-Year Longitudinal Cohort Study.” JAMA Neurol 75(3): 279–286.

Cugurra, A., T. Mamuladze, J. Rustenhoven, T. Dykstra, G. Beroshvili, Z. J. Greenberg, W. Baker, Z. Papadopoulos, A. Drieu, S. Blackburn, M. Kanamori, S. Brioschi, J. Herz, L. G. Schuettpelz, M. Colonna, I. Smirnov and J. Kipnis (2021). “Skull and vertebral bone marrow are myeloid cell reservoirs for the meninges and CNS parenchyma.” Science 373(6553).

Cusick, M. F., J. E. Libbey, D. C. Patel, D. J. Doty and R. S. Fujinami (2013). “Infiltrating macrophages are key to the development of seizures following virus infection.” J Virol 87(3): 1849–1860.

Deacon, R. M. (2006). “Assessing nest building in mice.” Nat Protoc 1(3): 1117–1119.

DePaula-Silva, A. B., F. L. Sonderegger, J. E. Libbey, D. J. Doty and R. S. Fujinami (2018). “The immune response to picornavirus infection and the effect of immune manipulation on acute seizures.” J Neurovirol 24(4): 464–477.

Dingledine, R., N. H. Varvel and F. E. Dudek (2014). “When and how do seizures kill neurons, and is cell death relevant to epileptogenesis?” Adv Exp Med Biol 813: 109–122.

Dingledine, R., N. H. Varvel, T. Ravizza and A. Vezzani (2024). Neuroinflammation in Epilepsy: Cellular and Molecular Mechanisms. Jasper’s Basic Mechanisms of the Epilepsies. J. L. Noebels, M. Avoli, M. A. Rogawski, A. Vezzani and A. V. Delgado-Escueta. New York: 611–632.

Dube, C. M., J. L. Zhou, M. Hamamura, Q. Zhao, A. Ring, J. Abrahams, K. McIntyre, O. Nalcioglu, T. Shatskih, T. Z. Baram and G. L. Holmes (2009). “Cognitive dysfunction after experimental febrile seizures.” Exp Neurol 215(1): 167–177.

Feng, L., M. Murugan, D. B. Bosco, Y. Liu, J. Peng, G. A. Worrell, H. L. Wang, L. E. Ta, J. R. Richardson, Y. Shen and L. J. Wu (2019). “Microglial proliferation and monocyte infiltration contribute to microgliosis following status epilepticus.” Glia 67(8): 1434–1448.

Franklin, K. a. P., G. (1996). “Mouse Brain in Stereotaxic Coordinates” Morgan Kaufmann, New York, NY.

Fu, Y., M. S. Yang, J. Jiang, T. Ganesh, E. Joe and R. Dingledine (2015). “EP2 Receptor Signaling Regulates Microglia Death.” Mol Pharmacol 88(1): 161–170.

Galanopoulou, A. S., M. Wong, D. Binder, A. L. Hartman, E. M. Powell, A. Roopra, R. Staba, A. Vezzani, B. Fureman, R. Dingledine, D. American Epilepsy Society /National Institute of Neurological and S. Stroke Epilepsy Benchmarks (2016). “2014 Epilepsy Benchmarks Area II: Prevent Epilepsy and Its Progression.” Epilepsy Curr 16(3): 187–191.

Goldman, A. M., W. C. LaFrance, Jr., T. Benke, M. Asato, D. Drane, A. Pack, T. Syed, R. Doss, S. Lhatoo, B. Fureman, R. Dingledine, D. American Epilepsy Society /National Institute of Neurological and S. Stroke Epilepsy Benchmark (2016). “2014 Epilepsy Benchmarks Area IV: Limit or Prevent Adverse Consequence of Seizures and Their Treatment Across The Lifespan.” Epilepsy Curr 16(3): 198–205.

Helmstaedter, C. (2007). “Cognitive outcome of status epilepticus in adults.” Epilepsia 48 Suppl 8: 85–90.

Heneka, M. T., W. M. van der Flier, F. Jessen, J. Hoozemanns, D. R. Thal, D. Boche, F. Brosseron, C. Teunissen, H. Zetterberg, A. H. Jacobs, P. Edison, A. Ramirez, C. Cruchaga, J. C. Lambert, A. R. Laza, J. V. Sanchez-Mut, A. Fischer, S. Castro-Gomez, T. D. Stein, L. Kleineidam, M. Wagner, J. J. Neher, C. Cunningham, S. K. Singhrao, M. Prinz, C. K. Glass, J. C. M. Schlachetzki, O. Butovsky, K. Kleemann, P. L. De Jaeger, H. Scheiblich, G. C. Brown, G. Landreth, M. Moutinho, J. Grutzendler, D. Gomez-Nicola, R. M. McManus, K. Andreasson, C. Ising, D. Karabag, D. J. Baker, S. A. Liddelow, A. Verkhratsky, M. Tansey, A. Monsonego, L. Aigner, G. Dorothee, K. A. Nave, M. Simons, G. Constantin, N. Rosenzweig, A. Pascual, G. C. Petzold, J. Kipnis, C. Venegas, M. Colonna, J. Walter, A. J. Tenner, M. K. O’Banion, J. R. Steinert, D. L. Feinstein, M. Sastre, K. Bhaskar, S. Hong, D. P. Schafer, T. Golde, R. M. Ransohoff, D. Morgan, J. Breitner, R. Mancuso and S. P. Riechers (2024). “Neuroinflammation in Alzheimer disease.” Nat Rev Immunol.

Hort, J., G. Brozek, P. Mares, M. Langmeier and V. Komarek (1999). “Cognitive functions after pilocarpine-induced status epilepticus: changes during silent period precede appearance of spontaneous recurrent seizures.” Epilepsia 40(9): 1177–1183.

Jiang, C., A. Caskurlu, T. Ganesh and R. Dingledine (2020). “Inhibition of the prostaglandin EP2 receptor prevents long-term cognitive impairment in a model of systemic inflammation.” Brain Behav Immun Health 8: 100132.

Jiang, J., T. Ganesh, Y. Du, Y. Quan, G. Serrano, M. Qui, I. Speigel, A. Rojas, N. Lelutiu and R. Dingledine (2012). “Small molecule antagonist reveals seizure-induced mediation of neuronal injury by prostaglandin E2 receptor subtype EP2.” Proc Natl Acad Sci U S A 109(8): 3149–3154.

Jiang, J., M. S. Yang, Y. Quan, P. Gueorguieva, T. Ganesh and R. Dingledine (2015). “Therapeutic window for cyclooxygenase-2 related anti-inflammatory therapy after status epilepticus.” Neurobiol Dis 76: 126–136.

Kanner, A. M. (2016). “Management of psychiatric and neurological comorbidities in epilepsy.” Nat Rev Neurol 12(2): 106–116.

Kantanen, A. M., M. Reinikainen, I. Parviainen and R. Kalviainen (2017). “Long-term outcome of refractory status epilepticus in adults: A retrospective population-based study.” Epilepsy Res 133: 13–21.

Kaufer, C., C. Chhatbar, S. Broer, I. Waltl, L. Ghita, I. Gerhauser, U. Kalinke and W. Loscher (2018). “Chemokine receptors CCR2 and CX3CR1 regulate viral encephalitis-induced hippocampal damage but not seizures.” Proc Natl Acad Sci U S A 115(38): E8929–E8938.

Kleen, J. K., R. C. Scott, P. P. Lenck-Santini and G. L. Holmes (2012). Cognitive and Behavioral Co-Morbidities of Epilepsy. Jasper’s Basic Mechanisms of the Epilepsies. J. L. Noebels, M. Avoli, M. A. Rogawski, R. W. Olsen and A. V. Delgado-Escueta. Bethesda (MD).

Leeman-Markowski, B. A. and S. C. Schachter (2016). “Treatment of Cognitive Deficits in Epilepsy.” Neurol Clin 34(1): 183–204.

Legriel, S., E. Azoulay, M. Resche-Rigon, V. Lemiale, B. Mourvillier, A. Kouatchet, G. Troche, M. Wolf, R. Galliot, G. Dessertaine, D. Combaux, F. Jacobs, P. Beuret, B. Megarbane, P. Carli, Y. Lambert, F. Bruneel and J. P. Bedos (2010). “Functional outcome after convulsive status epilepticus.” Crit Care Med 38(12): 2295–2303.

Levin, J. R., G. Serrano and R. Dingledine (2012). “Reduction in delayed mortality and subtle improvement in retrograde memory performance in pilocarpine-treated mice with conditional neuronal deletion of cyclooxygenase-2 gene.” Epilepsia 53(8): 1411–1420.

Lu, E., N. Pyatka, C. J. Burant and M. Sajatovic (2021). “Systematic Literature Review of Psychiatric Comorbidities in Adults with Epilepsy.” J Clin Neurol 17(2): 176–186.

Lu, M., M. Faure, A. Bergamasco, W. Spalding, A. Benitez, Y. Moride and M. Fournier (2020). “Epidemiology of status epilepticus in the United States: A systematic review.” Epilepsy Behav 112: 107459.

Meador, K. J. (2006). “Cognitive and memory effects of the new antiepileptic drugs.” Epilepsy Res 68(1): 63–67.

Minhas, P. S., A. Latif-Hernandez, M. R. McReynolds, A. S. Durairaj, Q. Wang, A. Rubin, A. U. Joshi, J. Q. He, E. Gauba, L. Liu, C. Wang, M. Linde, Y. Sugiura, P. K. Moon, R. Majeti, M. Suematsu, D. Mochly-Rosen, I. L. Weissman, F. M. Longo, J. D. Rabinowitz and K. I. Andreasson (2021). “Restoring metabolism of myeloid cells reverses cognitive decline in ageing.” Nature 590(7844): 122–128.

Mohammad, H., S. Sekar, Z. Wei, F. Moien-Afshari and C. Taghibiglou (2019). “Perampanel but Not Amantadine Prevents Behavioral Alterations and Epileptogenesis in Pilocarpine Rat Model of Status Epilepticus.” Mol Neurobiol 56(4): 2508–2523.

Mula, M., H. Coleman and S. J. Wilson (2022). “Neuropsychiatric and Cognitive Comorbidities in Epilepsy.” Continuum (Minneap Minn) 28(2): 457–482.

Parent, J. M. (2008). “Persistent hippocampal neurogenesis and epilepsy.” Epilepsia 49 Suppl 5: 1–2.

Parent, J. M., V. V. Valentin and D. H. Lowenstein (2002). “Prolonged seizures increase proliferating neuroblasts in the adult rat subventricular zone-olfactory bulb pathway.” J Neurosci 22(8): 3174–3188.

Park, S. P. and S. H. Kwon (2008). “Cognitive effects of antiepileptic drugs.” J Clin Neurol 4(3): 99–106.

Paudel, Y. N., M. F. Shaikh, S. Shah, Y. Kumari and I. Othman (2018). “Role of inflammation in epilepsy and neurobehavioral comorbidities: Implication for therapy.” Eur J Pharmacol 837: 145–155.

Power, K. N., A. Gramstad, N. E. Gilhus, K. O. Hufthammer and B. A. Engelsen (2018). “Cognitive dysfunction after generalized tonic-clonic status epilepticus in adults.” Acta Neurol Scand 137(4): 417–424.

Power, K. N., A. Gramstad, N. E. Gilhus, K. O. Hufthammer and B. A. Engelsen (2018). “Cognitive function after status epilepticus versus after multiple generalized tonic-clonic seizures.” Epilepsy Res 140: 39–45.

Quan, Y., J. Jiang and R. Dingledine (2013). “EP2 receptor signaling pathways regulate classical activation of microglia.” J Biol Chem 288(13): 9293–9302.

Rojas, A., R. Amaradhi, A. Banik, C. Jiang, J. Abreu-Melon, S. Wang, R. Dingledine and T. Ganesh (2021). “A Novel Second-Generation EP2 Receptor Antagonist Reduces Neuroinflammation and Gliosis After Status Epilepticus in Rats.” Neurotherapeutics 18(2): 1207–1225.

Rojas, A., T. Ganesh, N. Lelutiu, P. Gueorguieva and R. Dingledine (2015). “Inhibition of the prostaglandin EP2 receptor is neuroprotective and accelerates functional recovery in a rat model of organophosphorus induced status epilepticus.” Neuropharmacology 93: 15–27.

Rojas, A., T. Ganesh, Z. Manji, T. O’Neill and R. Dingledine (2016). “Inhibition of the prostaglandin E2 receptor EP2 prevents status epilepticus-induced deficits in the novel object recognition task in rats.” Neuropharmacology 110(Pt A): 419–430.

Rojas, A., J. Jiang, T. Ganesh, M. S. Yang, N. Lelutiu, P. Gueorguieva and R. Dingledine (2014). “Cyclooxygenase-2 in epilepsy.” Epilepsia 55(1): 17–25.

Rojas, A., H. S. McCarren, J. Wang, W. Wang, J. Abreu-Melon, S. Wang, J. H. McDonough and R. Dingledine (2021). “Comparison of neuropathology in rats following status epilepticus induced by diisopropylfluorophosphate and soman.” Neurotoxicology 83: 14–27.

Scharfman, H. E. and W. P. Gray (2007). “Relevance of seizure-induced neurogenesis in animal models of epilepsy to the etiology of temporal lobe epilepsy.” Epilepsia 48 Suppl 2(Suppl 2): 33–41.

Sharma, R., E. Chu, L. K. Dill, A. Shad, A. Zamani, T. J. O’Brien, P. M. Casillas-Espinosa, S. R. Shultz and B. D. Semple (2023). “Ccr2 Gene Ablation Does Not Influence Seizure Susceptibility, Tissue Damage, or Cellular Inflammation after Murine Pediatric Traumatic Brain Injury.” J Neurotrauma 40(3-4): 365–382.

Tian, D. S., J. Peng, M. Murugan, L. J. Feng, J. L. Liu, U. B. Eyo, L. J. Zhou, R. Mogilevsky, W. Wang and L. J. Wu (2017). “Chemokine CCL2-CCR2 Signaling Induces Neuronal Cell Death via STAT3 Activation and IL-1beta Production after Status Epilepticus.” J Neurosci 37(33): 7878–7892.

Varvel, N. H., R. Amaradhi, C. Espinosa-Garcia, S. Duddy, R. Franklin, A. Banik, C. Aleman-Ruiz, L. Blackmar-Raynolds, W. Wang, T. Honore, T. Ganesh and R. Dingledine (2023). “Preclinical development of an EP2 antagonist for post-seizure cognitive deficits.” Neuropharmacology 224: 109356.

Varvel, N. H., C. Espinosa-Garcia, S. Hunter-Chang, D. Chen, A. Biegel, A. Hsieh, L. Blackmer-Raynolds, T. Ganesh and R. Dingledine (2021). “Peripheral Myeloid Cell EP2 Activation Contributes to the Deleterious Consequences of Status Epilepticus.” J Neurosci 41(5): 1105–1117.

Varvel, N. H., J. J. Neher, A. Bosch, W. Wang, R. M. Ransohoff, R. J. Miller and R. Dingledine (2016). “Infiltrating monocytes promote brain inflammation and exacerbate neuronal damage after status epilepticus.” Proc Natl Acad Sci U S A 113(38): E5665–5674.

Waltl, I., C. Kaufer, S. Broer, C. Chhatbar, L. Ghita, I. Gerhauser, M. Anjum, U. Kalinke and W. Loscher (2018). “Macrophage depletion by liposome-encapsulated clodronate suppresses seizures but not hippocampal damage after acute viral encephalitis.” Neurobiol Dis 110: 192–205.

West, M. J. (1993). “New stereological methods for counting neurons.” Neurobiol Aging 14(4): 275–285.

Witt, J. A. and C. Helmstaedter (2015). “Cognition in the early stages of adult epilepsy.” Seizure 26: 65–68.

